# Small dendritic synapses enhance temporal coding in a model of cochlear nucleus bushy cells

**DOI:** 10.1101/2020.06.03.131516

**Authors:** Elisabeth Koert, Thomas Kuenzel

## Abstract

Spherical bushy cells (SBC) in the the anteroventral cochlear nucleus can improve the temporal precision of the auditory nerve spiking activity despite receiving sometimes only a single suprathreshold axosomatic input. The interaction with small dendritic inputs could provide a possible explanation for this phenomenon. In a compartment model of spherical bushy cells with a stylized or realistic three-dimensional representation of the bushy dendrite we explored this proposal. Phase-locked dendritic inputs caused both a tonic depolarization and a modulation of the SBC membrane potential at the frequency of the stimulus but for plausible model parameters do not cause output action potentials (AP). The tonic depolarization increased the excitability of the SBC model. The modulation of the membrane potential caused a phase-dependent increase in the efficacy of the main axosomatic input to cause output AP. These effects increased the rate and the temporal precision of output AP. Rate was mainly increased for stimulus frequencies at and below the characteristic frequency of the main input. Precision mostly increased for higher frequencies above about 1 kHz. Dendritic morphological parameters, biophysical parameters of the dendrite and the synaptic inputs and tonotopic parameters of the inputs all affected the impact of dendritic synapses. This suggested the possibility of fine tuning of the effect the dendritic inputs have for different coding demands or input frequency ranges. Excitatory dendritic inputs modulate the processing of the main input and are thus a plausible mechanism for the improvement of temporal precision in spherical bushy cells.

## Introduction

Spherical bushy cells (SBC) are monaural neurons in the anteroventral part of the mammalian cochlear nucleus (Bazwinsky et al., 2008). They receive narrow-band input (Blackburn and Sachs, 1989) from the auditory nerve (AN) via the endbulb of Held axosomatic terminals (Ryugo and Parks, 2003; Felmy and Künzel, 2014) and project their axons to binaural nuclei in the sound localization circuitry of the superior olivary nucleus (Cant and Benson, 2003). SBC maintain the temporal code contained in AN action potentials and have been reported to improve the precision of phase locking (Joris et al., 1994a,b; Young and Sachs, 2008; Kuenzel et al., 2011; Keine and Rübsamen, 2015; Wei et al., 2017) sometimes beyond what individual AN fibers are capable of (Versteegh et al., 2011). The mechanism for this is well understood in the case of numerous phase-locked subthreshold inputs: several events have to precisely coincide to cause an action potential (Rothman et al., 1993; Xu-Friedman and Regehr, 2005a) which causes a narrower distribution of output phase angles. This seemed to be the case for SBC with high characteristic frequencies and for globular bushy cells (Spirou et al., 2005). Enhancement of temporal precision by the coincidence of numerous inputs has also been successfully demonstrated in computer simulations of SBC (Rothman et al., 1993; Rothman and Young, 1996). For SBC that receive only a small number of supra-threshold inputs (Cao and Oertel, 2010) a different mechanism was proposed: here only the first active suprathreshold input caused an output action potential (Xu-Friedman and Regehr, 2005b). This biased and sharpened the distribution of temporal positions contained in the output towards early suprathreshold events. Given that endbulb of Held terminals were reported to be much less effective in vivo (Borst, 2010; Kuenzel et al., 2011), inhibitory inputs to SBC (Kuenzel et al., 2011; Campagnola and Manis, 2014; Nerlich et al., 2014; Keine and Rübsamen, 2015; Kuenzel et al., 2015; Keine et al., 2016) also play a major role in both scenarios by dynamically setting the action potential threshold.

Interestingly, in a subset of low-frequency SBC that often receive only a single, very strong endbulb of Held input, enhancement of temporal precision was also observed (Smith et al., 1993; Joris et al., 1994a; Kuenzel et al., 2011). This finding cannot be explained by the mechanisms discussed above. This conundrum was termed the spherical cell puzzle (Joris and Smith, 2008). What is the source of additional information for these low-frequency, single-input SBC? First it has to be noted that even these low-frequency SBC do not only receive a single synaptic input. In fact a wide array of smaller inputs from various sources converge on the dendritic structures of SBC (Kuenzel, 2019), some being AN fibers (Gómez-Nieto and Rubio, 2009) of the same characteristic frequency as the endbulb input. We hypothesize that these phase-coupled weak inputs act as a “hidden” augmentation to the main input and represent the extra information needed to enhance the temporal precision in these neurons. In-vivo studies by Keine and Rübsamen (2015) and Kuenzel et al. (2011) indeed showed that the strongest EPSP of a single endbulb showed best timing. This was also a required feature for the increase in temporal precision in one of our prior modeling studies (Kuenzel et al., 2015). Unfortunately there is little concrete physiological information available on small excitatory dendritic inputs (Cao and Oertel, 2010) and the dendritic properties of SBC (Oertel et al., 2008). Based on morphological data it is assumed that dendritic AN inputs to SBC are numerous but weak, bouton-like contacts (Gómez-Nieto and Rubio, 2009). At least some were reported to be additional axodendritic connections of axosomatic AN terminals (Ostapoff and Morest, 1991; Ryugo and Sento, 1991; Gómez-Nieto and Rubio, 2009).

We therefore designed the following modeling study with two objectives in mind: first we wanted to explore whether and how a number of phase-locked but weak dendritic inputs can shape the input-output relation of the main axosomatic input, especially with regard to temporal coding. And second, we sought to deduce and discuss the range of physiological parameters of both the dendritic inputs and the dendritic structures of SBC at which phase-locked dendritic inputs could help temporal processing. To achieve these goals we developed a compartment model of SBC closely based on physiological data and attached either a stylized or a realistic 3D-dendritic structure. We explored what the impact of dendritic inputs was on SBC subthreshold membrane potentials as well as their interaction with the endbulb of Held terminal. Our results indicated that small phase-locked dendritic inputs can improve the encoding of temporal information contained in the main axosomatic input and thus represent one likely piece of the spherical cell puzzle.

## Material and Methods

### Staining and 3D-reconstruction of gerbil spherical bushy cells

For N = 18 spherical bushy cells filled with biocytin during whole-cell recordings in acute gerbil brain slices we obtained morphological data. Recording techniques and the staining approach was described in detail elsewhere before (Goyer et al., 2016; Gillet et al., 2018; Gillet et al., 2020). Briefly, cells were recovered post-hoc by fixation of the slices in 4 % paraformaldhyde in 0.1 M phosphate buffer overnight. After washing with buffered saline solution containing 0.3 % Triton X-100, slices were incubated with Alexa-streptavidin conjugate (Thero Fischer Scientific) diluted 1:800 in 0.1 M phosphate buffer containing 0.1 % Triton X-100 and 1 % bovine serum albumine for 3 h at RT. Washed (TRIS-buffered saline containing 0.3 % Triton X-100) slices were then mounted in Fluoprep medium (bioMerieux) surrounded by a spacer frame (240 µm adhesive tapes; Grace Bio-Labs) between two glass coverslips.

SBC were visualized with a confocal laser-scanning microscope (Leica TCS SP2, Leica Microsystems) at high resolution. The number of images per stack was adjusted to assure the z-resolution was ≤ 0.5 µm. Cellular structures in these confocal stacks were reconstructed in 3D using the “Simple Neurite Tracer” plugin (Longair et al., 2011) for FIJI/ImageJ (Schindelin et al., 2012). Besides a description of each dendritic section as a hierarchic list of paths defined by points in x-, y- and z-coordinates, this included a diameter d for every coordinate. The diameter of each segment was derived by volume fitting the dendritic segment in the confocal image as described in Longair et al. (2011). We defined an arbitrary minimal diameter of 1 µm for dendritic segments. This was done because the volume fitting procedure, being based on the local fluorescence intensity of the image, produced numerous segments with diameters close to 0 µm. Somatic and axonal structures were reconstructed but not further analyzed. Dendritic structures were analyzed by counting the intersects of dendritic segments with spheres of increasing radii in 3D space (Sholl, 1953; Malinowski et al., 2019). Data describing both the 3D-shape as well as the identity and connectivity of all dendritic segments were exported from the FIJI-plugin and imported into our custom Python model code (see below).

### Compartment model of gerbil spherical bushy cells

Using NEURON (version 7.6.2) as a module in Python 2.7 running in Ubuntu Linux Server 18.04 we created a compartment model of gerbil SBC as previously published (Nerlich et al., 2014; Kuenzel et al., 2015; Goyer et al., 2016). The python code necessary to generate the figures of this manuscript is available for inspection as a git repository at https://github.com/thkupy/sbcnrnpy. The model consisted of a somatic section, to which an axon model (consisting of an initial segment and a stretch of passive axon representing the first internode) and either a stylized or realistic dendritic representation was attached. The stylized dendrite consisted of a proximal dendrite, to which a distal dendrite was attached (resulting in a “ball-and-stick with axon” model). In these experiments we defined classical geometry for these sections in NEURON. For the experiment using the realistic dendritic representation we created and connected appropriate numbers of sections based on our reconstructed SBC morphology and defined the geometry of each section using the pt3d-notation of NEURON. In the realistic dendrite model the dendritic path which connected to the soma section was defined as the proximal dendrite, all other dendritic sections were called distal. The geometry of axon, soma and stylized dendrite and the biophysical parameters of all sections can be found in Table 1. The Nernst- and reversal-potentials were: E_Na_ = 50 mV, E_K_ = −70 mV, E_H_ = −43 mV, E_leak_ = −65 mV. Axial resistance of all segments was set to 150 Ω·cm. All simulations were calculated at fixed temporal resolution of dt = 10 µs, for a temperature of 35 °C.

**Table 1.**
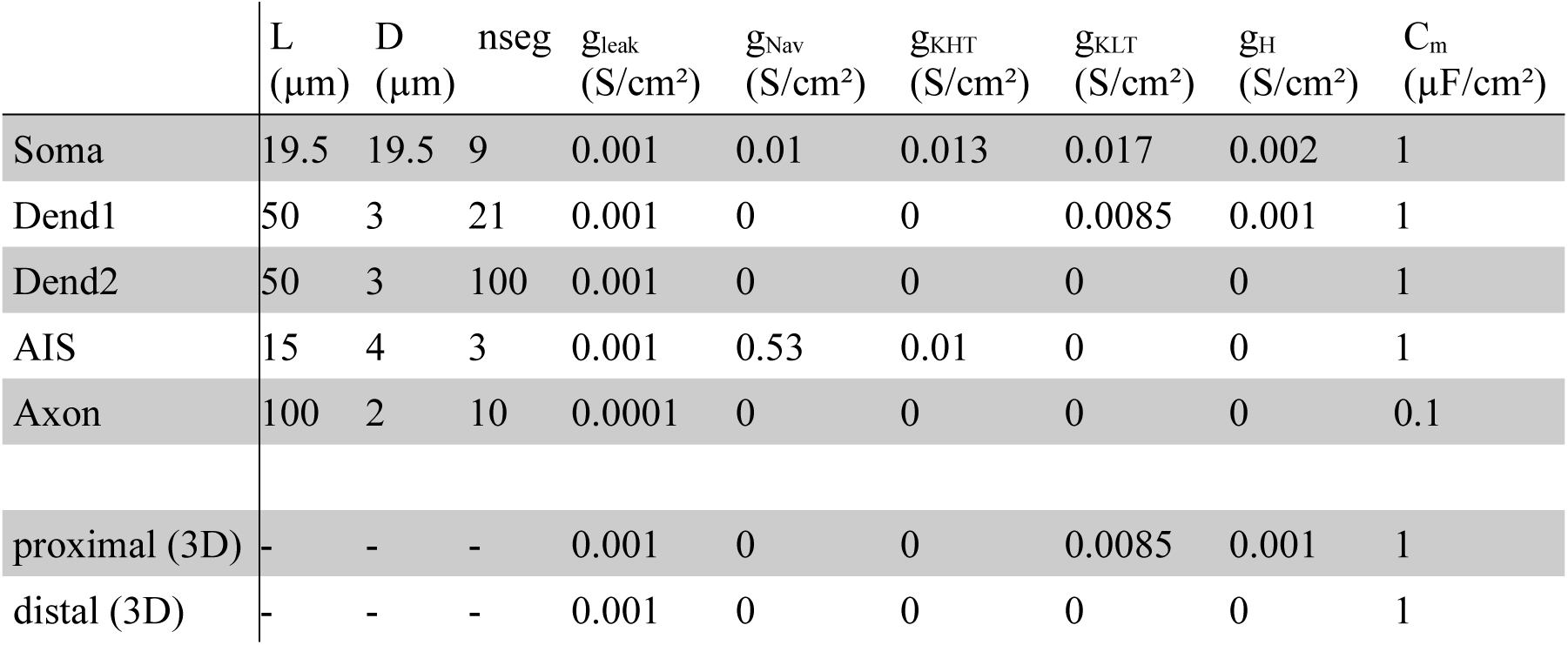
Morphological and biophysical parameters of the SBC compartment model.

We connected two classes of synaptic models to both the stylized and realistic SBC model. Dendritic synapses were per default modelled as N = 16 simple point synapses (*ExpSyn*) with a default synaptic conductance of g_syn_ = 0.5 nS, which were activated at specific times representing AN input events with the *NetCon* and *NetStim* mechanisms of NEURON. In the stylized model dendritic synapses were placed on the distal dendrite section only, with even spacing (spanning 5 % to 95 % of the section length). In the realistic dendrite model the dendritic connections were randomly chosen to predominantly (66 %) connect to the distal dendritic sections. Random dendritic positions were saved and reused for every simulation run (although a new random placement for every repetition produced, on average, the same results). Furthermore we attached a large conductance point source (modeled as a *gclamp* mechanism in NEURON) to the soma (at 50 % of the soma length) as a representation of the endbulb of Held input. Conductance traces for the endbulb of Held input were generated as described before (Nerlich et al., 2014; Kuenzel et al., 2015) to closely match the behavior of in-vivo endbulb of Held recordings. The synaptic conductance for the endbulb input was randomly varied for every input event (mean: 55 nS ± 11 nS). The reversal potential of all synapse models was set to Erev = 0 mV.

An overview of the model geometry and connectivity for the stylized dendrite model is shown in Fig. 1.

**Fig. 1.**
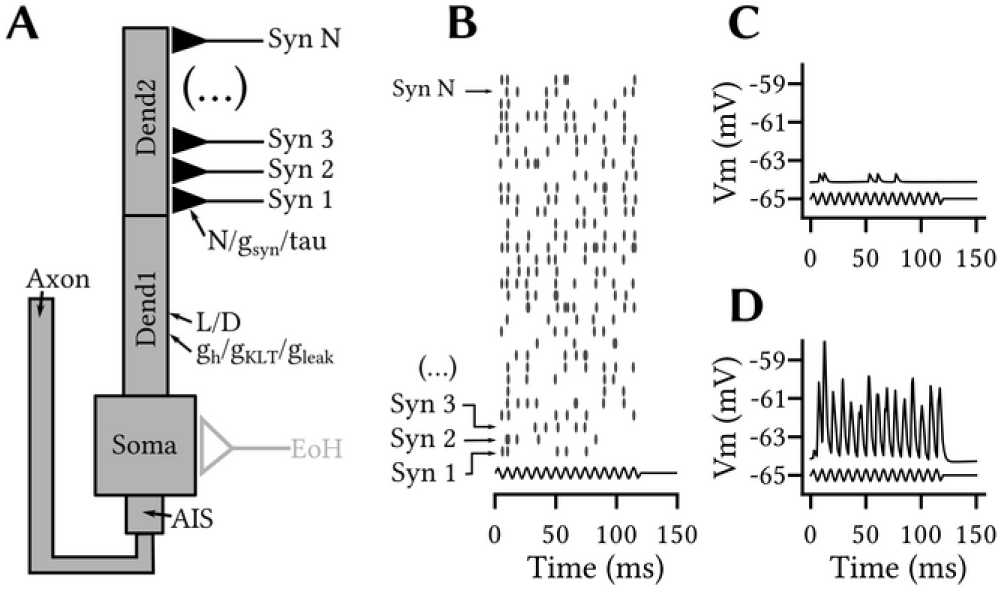
Phase-locked dendritic inputs caused both a tonic and a modulated subthreshold memberane response in the SBC model. *A:* Overview of the compartment model and parameters used in simulations. Note that for initial experiments the endbulb of Held (EoH) was not simulated. Dend1: primary dendrite section, Dend2: secondary dendrite section, AIS: axon initial segment, N: number of inputs, g_syn_: conductance of one dendritic synaptic input, tau: decay time-constant of dendritic synaptic inputs, L: length of primary dendrite section, D: diameter of primary dendrite section, Syn N: dendritic synapse number N, g_h_: hyperpolarization activated conductance, g_KLT_: low voltage activated potassium conductance, g_leak_: leak conductance. *B:* Spike times of N = 32 statistically independent simulated AN fibers upon 50 ms, 65 dB SPL, 125 Hz tone stimulation. Spike-times were used as activation times for the corresponding synaptic mechanisms. *C:* Membrane potential response at the soma of the SBC model upon activation of dendritic synapse #1 only. *D:* Membrane potential response at the soma of the SBC model upon activation of all N = 32 dendritic synapses. Note that synaptic events summated to cause both a modulation of the membrane potential at the stimulus frequency and an increase of the average membrane potential (tonic depolarization).

### Auditory nerve inputs

We modeled all inputs to the SBC model as sound driven auditory nerve spike arrival times generated by the well-known Zilany-model (Zilany et al., 2009; Zilany et al., 2014). All simulated sound responses were calculated at a temporal resolution of 10 µs (100 kHz) for medium spontaneous range fibers and the species-setting “cat”. Characteristic frequencies (CF) below 2.5 kHz were used in our simulations. For parallel activation of the dendritic and somatic synapse models we calculated AN sound driven responses as statistically independent AN fibers of the same CF. We used the Python module *cochlea* (Rudnicki et al., 2015) as a convenient wrapper for the Zilany-model.

### Data analysis and statistics

If not stated otherwise all results given in the paper are expressed as mean ± 1 standard deviation. Spike times in the simulated SBC membrane voltage responses were detected by thresholding the first derivative of the membrane potential, using the rapid falling flanks as distinct features of action potentials. The time of peak voltage of the action potential was then used as the spike time for further analysis. Failed endbulb of Held transmission was detected by the absence of an output spike in a time-window of 1 ms after a known endbulb input event. We calculated the precision of phase locking as the vector strength (Goldberg and Brown, 1969) of SBC output spike times. A confidence level of p < 0.001 (Rayleigh test) was used as a criterion for significance of phase locking (Fisher, 1993). Values of vector strength that failed the rayleigh criterion were set to 0. All data analysis and visualization was performed using the python modules *matplotlib* and *numpy*. For visualizing data as 2D contour-plots (*contourf*) mild smoothing was routinely applied to the data using a gaussian filter (σ = 0.5).

## Results

### Phase-locked excitatory dendritic inputs cause subthreshold membrane potential oscillations

We first analyzed the effect of small excitatory inputs to the model SBC dendrite without any main endbulb input (Fig. 1A). Dendritic inputs were driven by independent auditory nerve spike trains calculated for the same characteristic frequency (Fig. 1B). At the given inputs frequency and level used in this sparse low-frequency example (125 Hz, 60 dB SPL) a simulated AN fiber with a characteristic frequency (CF) of 250 Hz responded with 55 ± 16 AP/s and had a vector strength of 0.78 ± 0.1.

Activity of a single, weak (g_syn_ = 0.5 nS) dendritic input elicited small (0.41 mV) EPSP at the soma (Fig. 1C). The combined activity of a higher number (N = 32, g_total_syn_ = 16 nS) of dendritic inputs summated to sharp, albeit subthreshold, fluctuations of the membrane potential at the stimulus frequency (Fig. 1D). The amplitude of these fluctuations was 5.4 mV in this example. Furthermore, in addition to the phase-locked oscillations the summating dendritic EPSP also caused a small tonic depolarization of the resting membrane potential (V_rest_ = −64.5 mV without inputs, V_rest_ = −62.9 mV with inputs) during the simulated stimulus presentation. In the following experiments we wanted to better understand how the properties of the dendritic tree and the dendritic synaptic inputs shaped these membrane potential responses. Since actual physiological parameters of the SBC dendrite (ionic conductances, number and conductance of dendritic synapses etc.) are not readily available, we varied several parameters over a plausible range to derive their impact.

For this we averaged the SBC membrane potential over the stimulus cycle and quantified the mean membrane potential and the amount of membrane potential modulation. Both parameters strongly depended on the length (Fig. 2A1-B2) and the diameter (Fig. 2C1-D2) of the primary dendrite. While V_m_ depolarization and modulation fell exponentially with dendritic length (Fig. 2A1, A2), depolarization and modulation were maximal for a dendritic diameter of 3.3 µm (Fig. 2C1,C2) and were reduced for both smaller and wider diameters. The latter can be mostly explained by the increased leak conductance in the longer dendritic segment. To illustrate this point we again calculated these simulations for the different dendritic morphologies but kept the total leak conductance of the dendritic segment fixed (dashed lines in Fig. 2A1-C2). In this condition, V_m_ modulation still steeply drops with increasing length (Fig. 2A2) and decreasing diameters below about 3 µm (Fig. 2D2). Over all (Fig. 2E) our model showed that the effects of the dendritic inputs on V_m_ depolarization (Fig. 2E1) and V_m_ modulation (Fig. 2E2) mostly depended on the length and, given a minimum value of about 3 µm, to a lesser degree on the diameter of the primary dendrite. We concluded that in order to maximize the effect of the phase-locked dendritic inputs, primary dendrites of SBC should be as short as possible and thicker than 3 µm.

**Fig. 2.**
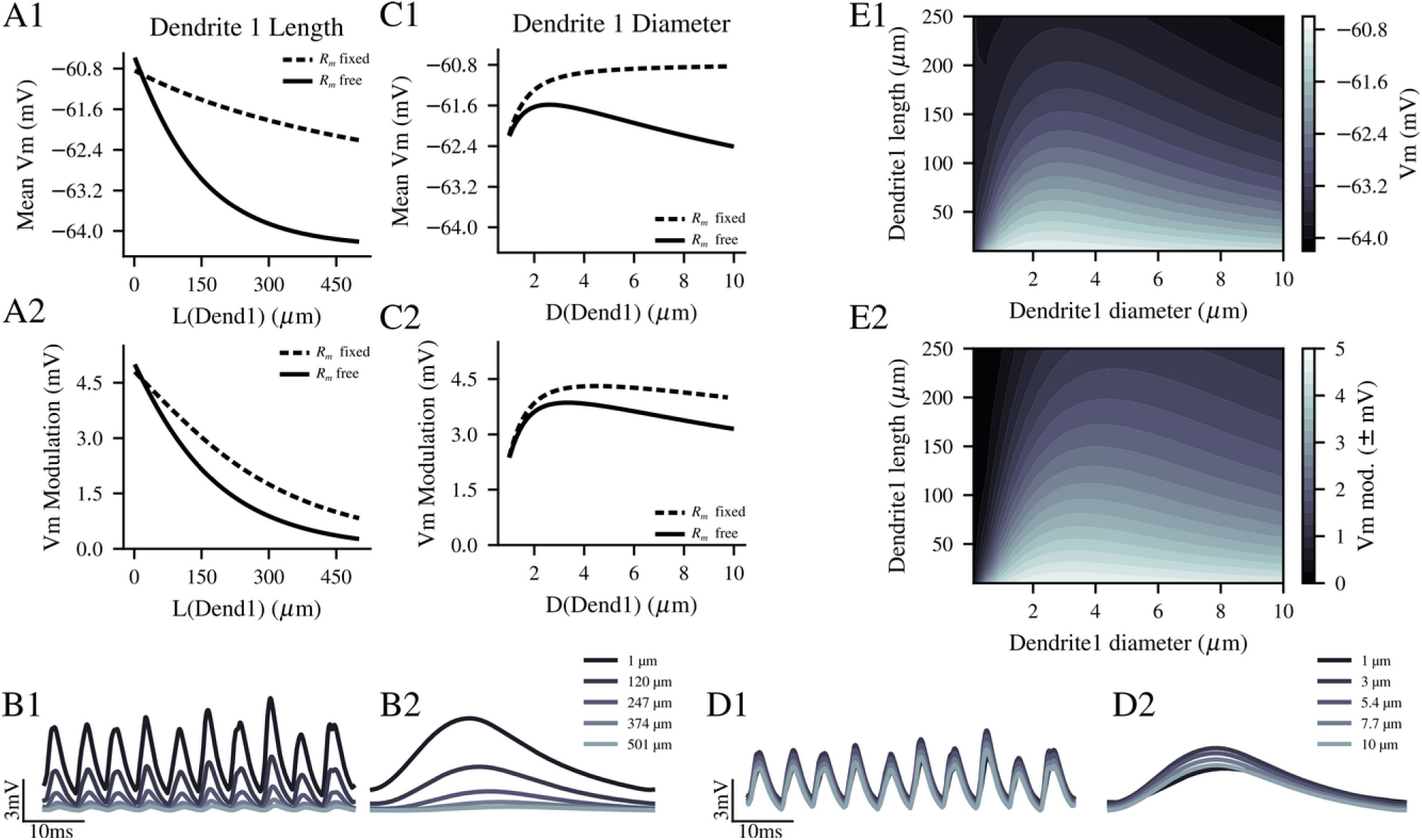
Morphology of the primary dendrite section influenced tonic and modulated subthreshold membrane responses of the model SBC caused by phase-locked dendritic inputs. *A:* Mean membrane potential (*A1*) and mean modulation amplitude (*A2*) were reduced with increasing length of the primary dendrite section L(Dend1), when D(Dend1) = 3 µm. Solid line: model with free total membrane resistance R_m_. In this model, R_m_ increased with increasing L, as more membrane area is added. Dashed line: model with fixed total membrane resistance R_m_. In this model, R_m_ was kept constant by reducing the specific leak conductance as the membrane area changed. For these and the following plot a minimum of 64 conditions were tested and at least 2 s of stimulus presentation was simulated per condition. *B:* Example traces (*B1*) and membrane potential averaged over the stimulus cycles (*B2*) for different L(Dend1). In *B1*, ten cycles of the stimulus are shown from 5 different example conditions. Linecolor in *B1* and *B2* indicates example values from low (dark) to high (bright). *C:* Mean membrane potential (*C1*) and mean modulation amplitude (*C2*) were maximal for dendrites of 3.3 µm and gradually reduced for diameters D(Dend1) above and below this value. Dendritic length was constant, L(Dend1) = 50 µm. *D:* Example traces (*D1*) and membrane potential averaged over the stimulus cycles (*D2*) for different D(Dend1). Presentation as in *B. E:* Contour-plots showing the mean membrane potential (*E1*) and mean modulation amplitude (*E2*) for 625 (25 x 25) combinations of L and D. Both parameters mostly depended on L(Dend1) with a specific slope that depended on D(Dend1).

Very little is known about the number of dendritic auditory nerve terminals of SBC (Gómez-Nieto and Rubio, 2009), and there is no good data on the synaptic conductance of these contacts (Cao and Oertel, 2010; maybe see). We thus wanted to explore the effect of a wide range of synapse numbers and total synaptic conductance on the membrane potential of the SBC (Fig. 3). We found that both the tonic depolarization and the membrane potential modulation did not depend on the actual number of inputs, above a low number of inputs (Fig. 3A1-B2). The depolarized resting membrane potential (RMP) converges to a value of −61.6 mV for N > 15 inputs, the RMP modulation reaches 3.84 mV above N > 23 inputs. In contrast to this, both the tonic depolarization and the RMP modulation monotonically increased with total synaptic conductance (Fig. 3C1-D2). Over a wide range of parameters (Fig. 3E), the influence of phase-locked synaptic inputs on SBC membrane potential almost exclusively depended on the total conductance. Only for very low numbers of inputs (N < 8) at high total conductances above 37 nS (Fig. 3E3) action potentials were elicited by dendritic inputs without endbulb of Held input. We concluded that a wide combination of the number and total conductance of dendritic auditory nerve terminals will only act on the subthreshold membrane potential of the SBC rather than directly cause output action potentials.

**Fig. 3.**
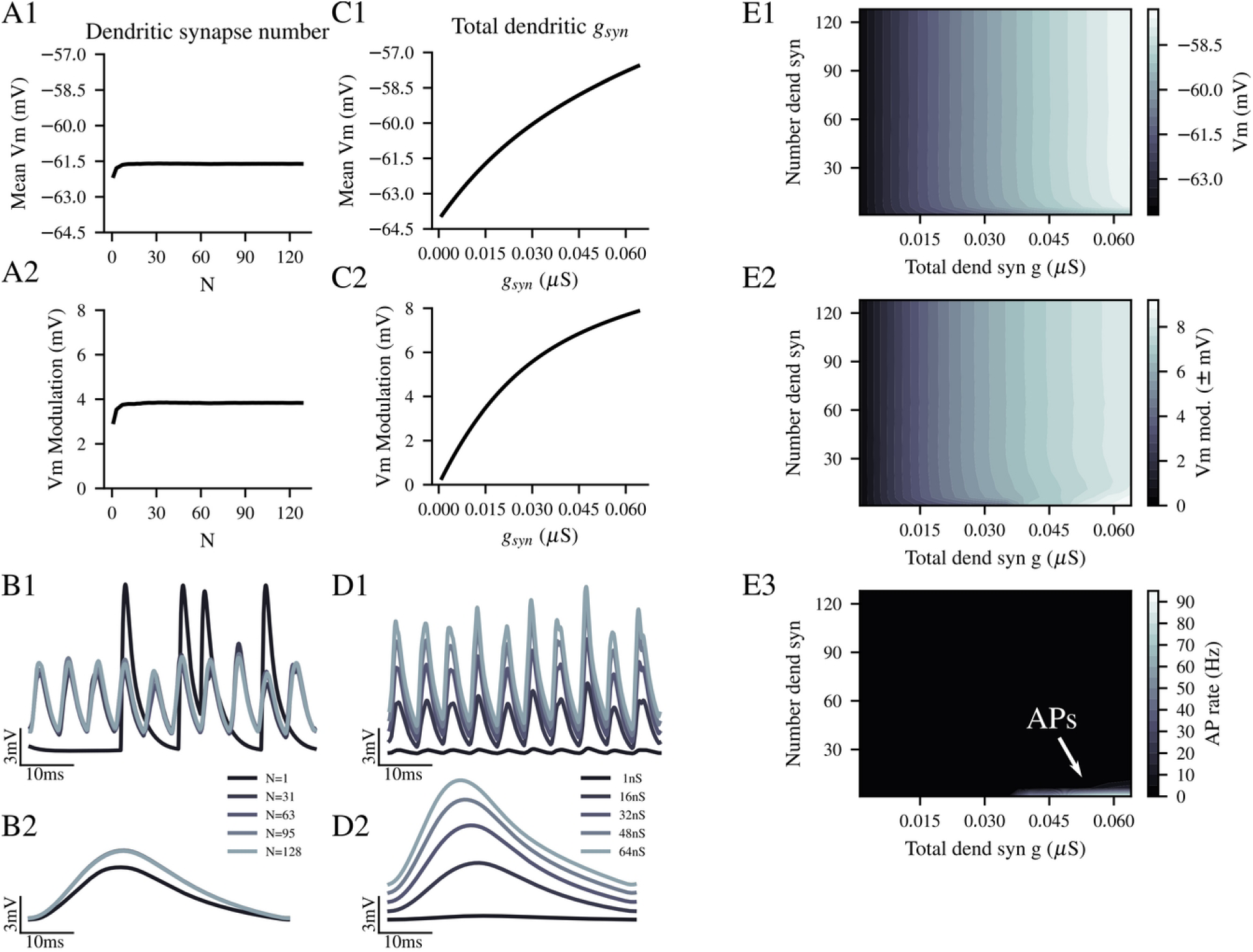
Effect of phase-locked dendritic inputs on tonic and modulated subthreshold SBC membrane responses was only weakly influenced by dendritic synapse number but strongly by total dendritic conductance. *A:* Mean membrane potential (*A1*) and mean modulation amplitude (*A2*) were not influenced by number of dendritic synapses N above N = 15 (*A1*) or N = 23 (*A2*). Total dendritic synaptic conductance was constant (g_syn_ = 16 nS). Below these low values parameters steeply declined with lower N. *B:* Example data for result shown in *A*, presentation as in Fig. 1B. *C:* Total dendritic synaptic conductance g_syn_ had a strong influence on mean membrane potential (*C1*) and mean modulation amplitude (*C2*). With increasing g_syn_ of N = 16 synapses, both parameters monotonically increased. *D:* Example data for results shown in *C*, presentation as in Fig. 1B. *E:* Contour-plots showing the mean membrane potential (*E1*) and mean modulation amplitude (*E2*) for 625 (25 x 25) combinations of N and g_syn_. Effects mainly depended on g_syn_. The number of action potentials triggered by dendritic inputs per condition is shown in the contour plot in *E3*.

Next, we wanted to explore the impact of the relation between the input cycle duration and the decay time-constant of the dendritic synaptic conductance on the SBC membrane potential (Fig. 4). One can assume that the synaptic currents elicited by glutamatergic auditory nerve terminals in the dendrites of SBC will have kinetics comparable to endbulb glutamatergic currents, but direct measurements are unavailable. Furthermore, it is obvious that the cycle duration of the phase-locked inputs will have a strong influence on the effect of summating synaptic potentials. With increasing decay time constant in response to 200 Hz input rate, the tonic depolarization of the RMP strongly increased (Fig. 4A1,B), as the synaptic events summated more efficiently. At the default value of τ_decay_ = 2 ms V_m_ was −61.5 mV, at τ_decay_ = 20 ms V_m_ was as depolarized as −53.3 mV. The amplitude of the RMP modulation, however, with increasing τ_decay_ rose steeply at first to maximum of 3.87 mV at τ_decay_ = 1.77 ms (Fig. 4A2,B). RMP modulation then gradually declined with increasing τ_decay_ to a value of 0.94 mV at the highest τ_decay_ tested (20 ms). Interestingly, due to the phase-locked nature of the inputs there was a small modulation at the frequency of the simulated sound stimulus visible in the membrane potential, even at these very long synaptic decay time-constants (Fig. 4B). With increasing cycle duration of the input at a fixed τ_decay_ = 2 ms, the tonic V_m_ depolarization slightly reduced (Fig. 4C1,D) from Vm = −60.5 mV (interval = 2 ms / 500 Hz) to Vm = −62.2 mV (at IPI = 7.5 ms / 133 Hz). The amplitude of Vm modulation converged to a value of 4.2 mV (at IPI = 7.5 ms / 133 Hz), starting from 1.4 mV (at IPI = 7.5 ms / 133 Hz). When we varied both the τ_decay_ and the IPI of the inputs (Fig. 4E1,E2) it became obvious that the contour lines in the 2d-plot ran mostly diagonally. This means that in order to ensure a constant amount of V_m_ depolarization (Fig. 4E1) or amplitude of V_m_ modulation (Fig. 4E2) over a wide range of mean IPI, the τ_decay_ of the inputs must be increased or decreased in accordance with the IPI. We thus concluded that, in order to cause both significant tonic and modulated effects on RMP, the dendritic auditory nerve terminals should have rapid decay time-constants of τ_decay_ < 5 ms. Furthermore the τ_decay_ of the dendritic synaptic inputs could provide a parameter to optimally tune the dendritic inputs to a specific range of input frequencies.

**Fig. 4.**
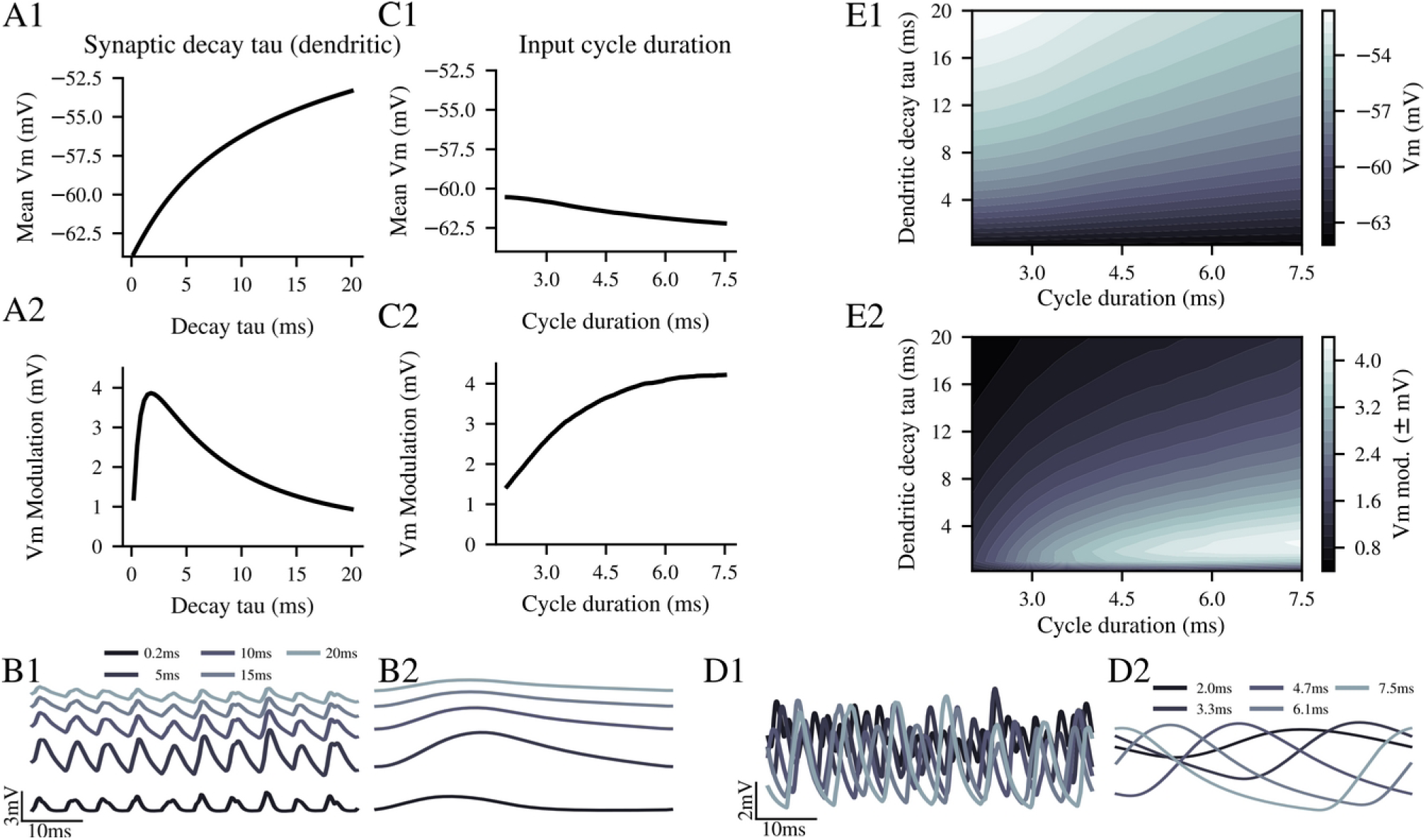
Temporal characteristics of inputs and dendritic synapses inversely affected tonic and modulated subthreshold membrane responses of the model SBC. *A:* Mean membrane potential (*A1*) and mean modulation amplitude (*A2*) were differentially affected by the decay time-constant of the dendritic synaptic conductance. Input frequency was 200 Hz. Mean membrane potential monotonically increased with decay tau, but mean modulation amplitude was maximal for tau = 1.77 ms and declined steeply for values above and below. *B:* Example data for result shown in *A*, presentation as in Fig. 1B. *C:* Input frequency, expressed here as duration of one cycle of the stimulus, weakly influenced mean membrane potential (*C1*). Shorter cycle duration / higher frequencies caused slightly higher tonic depolarization (for a synaptic decay time constant of tau = 2 ms). Mean modulation amplitude (*C2*) however increased strongly with increasing cycle duration. *D:* Example data for results shown in *C*, presentation as in Fig. 1B. *E:* Contour-plots showing the mean membrane potential (*E1*) and mean modulation amplitude (*E2*) for 625 (25 × 25) combinations of dendritic decay tau and cycle duration. The greatest variation of mean membrane potentials with decay tau occured for short cycle durations. Modulation amplitudes were greatest for a limited range of short decay time constants and long cycle durations.

It was reported (Oertel et al., 2008) that primary dendrites of SBC express both low-voltage activated potassium (g_KLT_) and hyperpolarization-activated cation conductance (g_H_). However, information about the actual density of either these voltage activated conductances or passive leak conductance (g _leak_) of the dendritic compartment were not available. We thus varied g_KLT_, g_H_ and g_leak_ over a range of plausible values and evaluated the impact of these parameters on the V_m_ depolarization and modulation by dendritic inputs (Figure 5). We found, that the amount of tonic V_m_ depolarization is reduced with increasing g_KLT_ (Fig. 5A1,B), while the amplitude of V_m_ modulation was largely unaffected (Fig. 5A2,B). In contrast to this, tonic depolarization slightly increased with increasing g_H_ (Fig. 5C1,D) without strong effects on the V_m_ modulation (Fig. 5C2,D). Thus the voltage-activated conductances that are coexpressed in SBC dendrites have opposing effects on the tonic depolarization but both leave the rapid, cycle-by-cycle modulation mostly unaffected. In contrast to this, increasing the passive leak conductance shunts both the tonic V_m_ depolarization (Fig. 5E1,F) and the V_m_ modulation (Fig. 5E2,F). Varying both g_KLT_ and g_H_ revealed (Fig. 5G1) that these conductances linearly counteracted each other in affecting the V_m_ depolarization. Thus, when both conductances increased roughly in a 2:1 ratio, the amount of V_m_ depolarization remained constant. On the other hand, g_KLT_ and g_H_ both had a small reducing effect on the amplitude of V_m_ modulation. This effect grew additively when both conductances increased (Fig. 5G2).

**Fig. 5.**
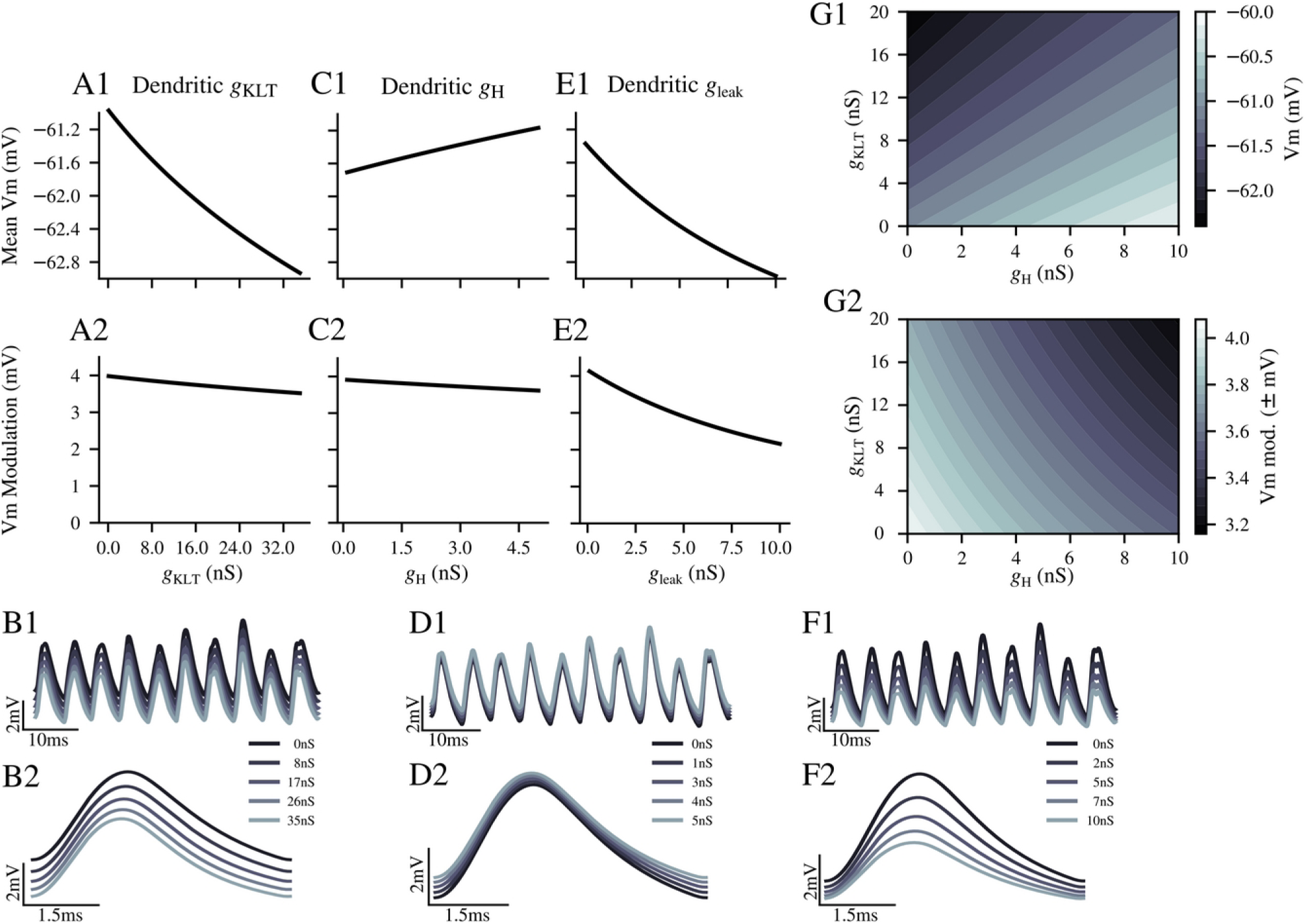
Passive and voltage-activated and dendritic conductances differentially affect tonic and modulated subthreshold membrane responses of the model SBC. *A:* Mean membrane potential (*A1*) was strongly reduced by low-voltage activated potassium conductance g_KLT_ but mean modulation amplitudes (*A2*) were hardly affected. *B:* Example data for result shown in *A*, presentation as in Fig. 1B. *C:* Mean membrane potential (*C1*) increased with increasing dendritic hyperpolarization-activated conductance g_h_. However, g_h_ had only very minor influence on mean modulation amplitudes (*C2*). *D:* Example data for result shown in *C*, presentation as in Fig. 1B. *E:* Both the mean membrane potential (*E1*) and mean modulation amplitude (*E2*) were strongly reduced by increasing dendritic passive leak conductance g_leak_. *F:* Example data for result shown in *E*, presentation as in Fig. 1B. *G:* Contour-plots showing the mean membrane potential (*G1*) and mean modulation amplitude (*G2*) for 625 (25 × 25) combinations of g_h_ and g_KLT_. G_h_ and g_KLT_ inversely affect the subthreshold membrane responses of the model SBC.

Overall our model results agree with the idea that SBC dendrites could be finely tuned to either emphasize the overall input level (tonic V_m_ depolarization) or the phase-locked nature (dynamic V_m_ modulation) of the dendritic synaptic inputs by the relative strength of g_KLT_, g_H_ and g_leak_ expressed in the dendritic compartment.

### Subthreshold membrane potential oscillations caused by dendritic inputs increase spike probability and timing precision of main endbulb inputs

Our model showed both a tonic and dynamic influence of dendritic inputs on subthreshold SBC V_m_. In the next step we assessed how the dendritic inputs could interact with the endbulb of Held axosomatic input. We thus modified the model to include a strong, somatically localized excitatory synaptic model that was driven by independent auditory nerve spike trains calculated for the same characteristic frequency as for the dendritic synapses (Fig. 6A). In the example shown in Fig. 6 (CF = 1000 Hz, @1000 Hz / 65 dB SPL) this resulted in simulated AN activity of 211 events/s driven rate and 55 events/s spontaneous rate. No short-term synaptic dynamics were simulated, however a stochastic variation of the endbulb EPSG amplitudes was implemented. This was shown before (Kuenzel et al., 2011; Nerlich et al., 2014; Kuenzel et al., 2015) to fit the in-vivo properties of this synapse well. The model SBC, without dendritic inputs, produced in this example a driven response of 180 AP/s and 50 AP/s spontaneous activity (Fig. 6B). Since the distribution of EPSG amplitudes (55 ± 11 nS) partially straddles the AP threshold of the model SBC, a moderate amount of failures occurred during simulated sound stimuli responses (31 failures/s) and during spontaneous activity (5 failures/s), as was reported for SBC from in vivo studies (Kuenzel et al., 2011; Keine et al., 2017). The auditory nerve inputs and therefore the output spikes of the model SBC showed robust phase locking (Fig. 6C) to the input fine-structure (AN inputs: VS = 0.80; SBC output without dendritic synapses: VS = 0.71). Interaction of the axosomatic input with dendritic inputs resulted in a higher output rate of the SBC model of 198 AP/s (Fig. 6D). The higher output rate was caused by a lower driven failure rate of only 13 failures/s. Spontaneous failures were identical for the two example conditions. Furthermore, the resulting output spikes were phase-locked more precisely to the input frequency (Fig. 6E), now showing a vector strength of 0.75. The improved temporal precision was accompanied by an advance of the preferred phase (φ = 0.23 cycles vs. φ = 0.28 cycles). The effects of dendritic inputs on temporal precision will be explored in greater detail in the following section. Inititally, we elucidated the mechanism by which the interaction of dendritic and somatic inputs increased the output spike rate of the model SBC.

**Fig. 6.**
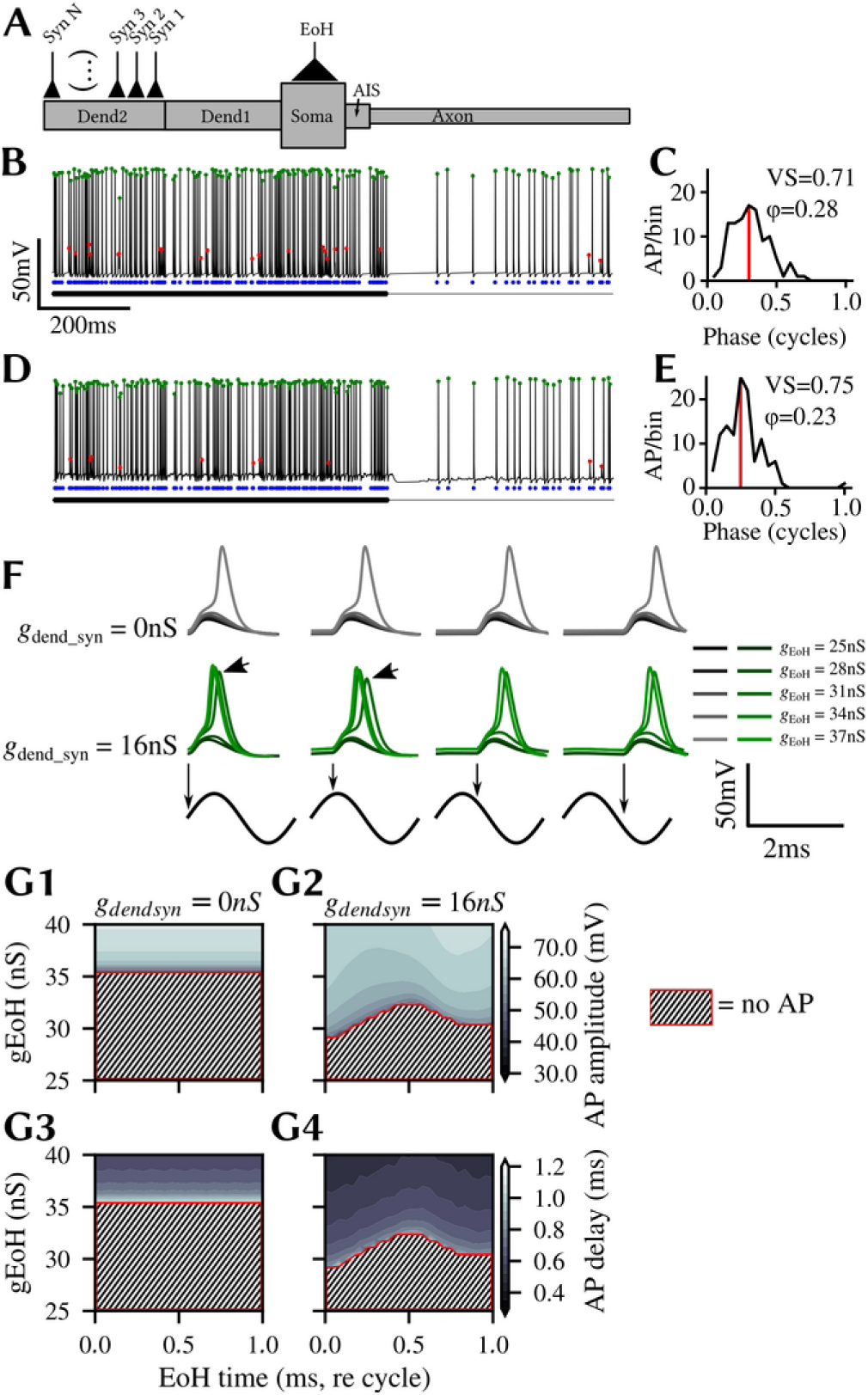
Dendritic inputs enhance endbulb of Held transmission efficacy. *A:* Schematic of the model used in the following experiments. The endbulb of Held axosomatic input is modeled as a conductance point source connected to the soma. *B:* Example trace of membrane potential responses of the SBC model with endbulb of Held input but without dendritic inputs upon simulated sound stimulation (1000 Hz, 65 dB SPL, CF = 1000 Hz). Successful endbulb to SBC transmission events, i.e. action potentials, marked with green dots. Failed transmissions, i.e. excitatory postsynaptic potentials, marked with red dots. Blue dots show timing of endbulb input events. All markers refer to the same time axis. Thick black line shows duration of simulated stimulus presentation. *C:* Cycle histogram of spike times relative to the input cycle for the SBC output spikes in *B*. Red vertical line shows mean phase angle of the events (preferred phase φ). *D:* Example trace of membrane potential responses of the SBC model with endbulb of Held input with N = 32, g_syn_ = 16 nS dendritic inputs upon simulated sound stimulation (1000 Hz, 65 dB SPL, CF = 1000 Hz). Presentation as in B. Note the lower amount of failed transmissions. *E:* Cycle histogram of spike times relative to the input cycle for the SBC output spikes in *D.* Presentation as in *C*. Note the higher amount of events, the higher vector strength and the phase advance. *F:* Phase relation of endbuld of Held input events and the membrane potential modulation caused by the dendritic inputs determines the efficacy of the endbulb of Held input. Examplary single activations of the endbulb of Held synapse at 25 (5 × 5) different combinations of peak conductance values and EoH activation times in relation to the stimulus cycle (1 s for a 1000Hz stimulus). Upper row, no dendritic inputs. Shades of grey indicate different endbulb peak conductance values. Middle row, with 16 nS dendritic synaptic inputs, shades of green indicate different endbulb peak conductance values. Bottom row, representation of the stimulus waveform. Vertical arrow marks temporal position of endbulb activation. *G:* Contour plots showing AP amplitude (*G1, G2*) and endbulb activation to peak AP delay (*G3, G4*) for 625 (25 x 25) combinations of EoH activation times and peak conductance without (*G1, G3*) and with (*G1, G4*) dendritic inputs. Hashed area with red boundary indicates conditions in which no AP was elicited.

For this we simulated single endbulb events of varying EPSG amplitude at specific fixed positions on the stimulus cycle, in order to estimate the conductance threshold at different phases of the dynamic V_m_ modulation caused by the dendritic inputs (Fig. 6F). Unsurprisingly, in the condition without dendritic inputs (Fig. 6F, upper row) the phase at which endbulb inputs occured had no effect on the conductance threshold. Below g_EoH_ = 37nS no action potentials were elicited by the endbulb events regardless of phase. In the condition with dendritic inputs events that occurred in the rising phase had a smaller conductance threshold (g_EoH_ = 31 nS) than events that occurred later in the cycle (Fig. 6F, middle row). Furthermore, on average the conductance to elicit an SBC action potential was lower in the dendritic input condition (g_EoH_ = 34 nS). We then quantified the results from the exemplary simulations shown in Fig. 6F by measuring the amplitude and delay of AP for a number of different EPSG amplitudes and phases (Fig. 6G). It was clear that in the model without dendritic inputs endbulb conductance threshold was not phase dependent (red line in Fig. 6G1/G3) and AP amplitude (Fig. 6G1) and delay (Fig. 6G3) only depended on g_EoH_. In the condition with dendritic inputs the phase dependence of the conductance threshold was evident (red line in Fig. 6G2/G4) resulting in a phase and g_EoH_ dependence of AP amplitude (Fig. 6G2) and delay (Fig. 6G4). Furthermore the dependence of AP delay on g_EoH_ at a given phase of the cycle was steeper, thus resulting in overall shorter AP delays. This explained the advance of preferred phase we observed (Fig. 6E). Taken together the quantification presented here suggested that dendritic inputs both tonically and dynamically reduced the conductance threshold of the endbulb, resulting in a phase-dependent improvement of the endbulb transmission efficacy.

### Frequency tuning of the interaction of dendritic inputs and main endbulb inputs

Since we found that the interaction of the dendritic inputs with the main endbulb inputs had an effect on temporal processing of the SBC model, we simulated auditory responses over a wide range of input stimulus frequencies and constructed frequency-response areas for the SBC model with and without dendritic inputs. As mentioned before, simulations were run with identical random seeds for each stimulus condition, to isolate the effect of the added dendritic synaptic conductance in the results. Figure 7A/B shows the primary-like response of the SBC model without and with the dendritic inputs. The overall shape and monotonically increasing characteristic of the response was unchanged. This was cleary evident by the 80 AP/s contour demarcating the responses clearly above spontaneous firing. However, the maximal response was increased in the condition with dendritic inputs, as was best shown by the larger extent of the area demarcated by the 170 AP/s contour (Fig. 7A/B). We plotted the difference between both conditions in the same coordiate system (Fig. 7C). It became obvious that the strongest increase in response rate caused by the dendritic inputs was at and below the CF of the simulated SBC. When we compared the effects of dendritic inputs on temporal aspects of the response, a different picture emerged (Fig. 7D-I). Precision of phase locking, expressed as VS and further illustrated in Fig. 7D/E with a VS = 0.6 contour, increased for frequencies at and above CF but remained largely unaffected in the low-frequency tail of the response (Fig. 7F). Indeed, the largest improvements of temporal precision occured within half an octave above the CF of the SBC for higher sound pressure levels (Fig. 7F). The accompanying advance of the preferred phase (Fig. 7G-I) showed a similar frequency dependence: it was greatest for frequencies at and above CF and weak below CF.

**Fig. 7.**
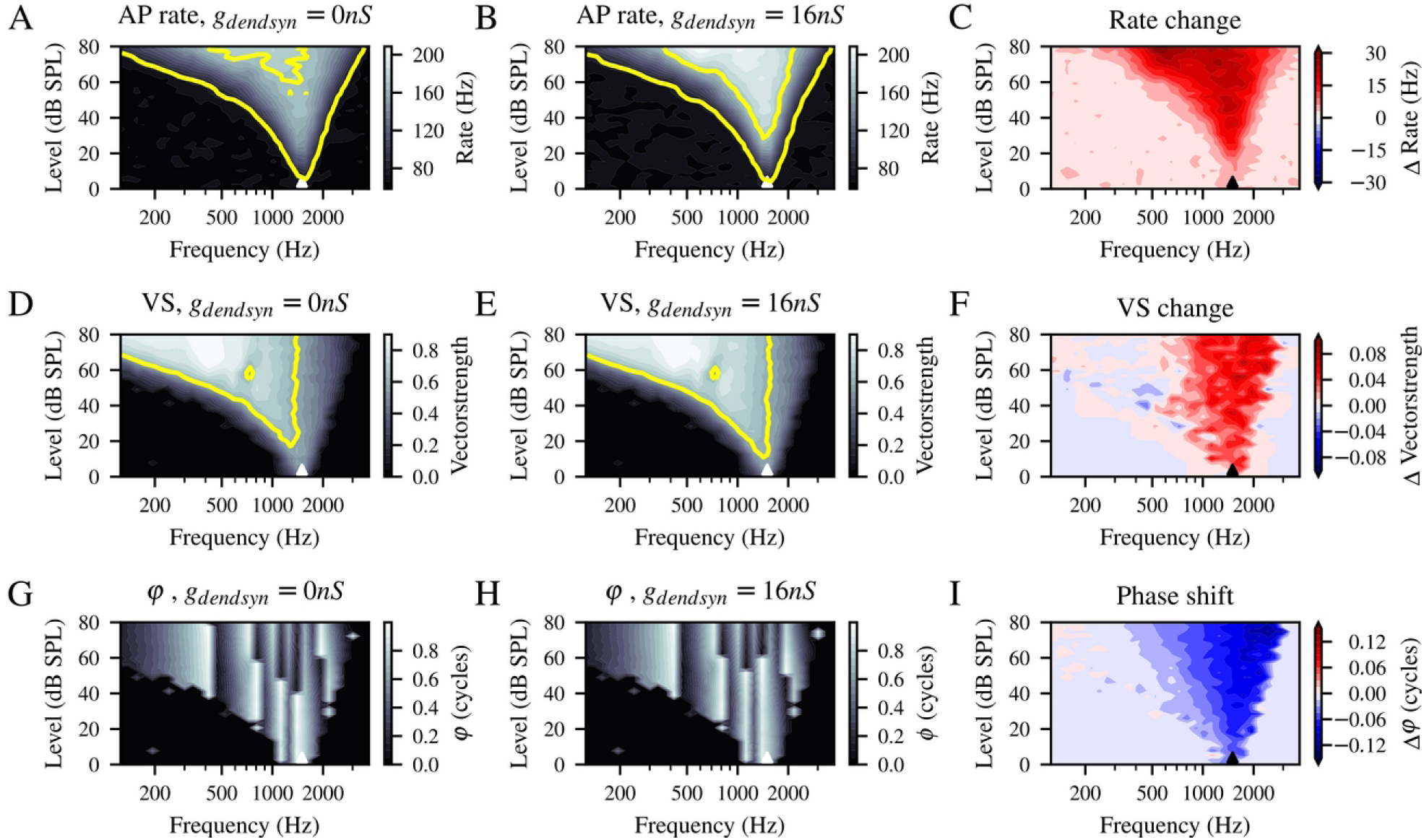
Dendritic inputs differentially affect SBC response rates and temporal coding dependent on stimulus condition. *A:* Contour plot of the frequency-response area of the SBC model without dendritic inputs, i.e. the output AP rate upon 1024 combinations of 32 different stimulus frequencies and 32 different stimulus sound pressure levels. 5 s of stimulus presentation were simulated per condition. CF = 1500 Hz. Yellow contours represent 80 Hz and 170 Hz output rate. *B:* Contour plot of the frequency-response area of the SBC model with N =32, gsyn = 16 nS dendritic inputs. Presentation as in A. Note the higher response rates visualized by the greater area demarcated by the 170 Hz contour. *C:* Difference between the data shown in A and B quantifies the effect of the dendritic inputs. Note the different color scheme: white signifies no difference, increasing intensity of red (blue) signifies increasing positive (negative) difference. Highest differences at and below CF. *D,E,F:* Contour plot of the temporal precision of the output AP quantified as vector strength without (*D*) and with (*E*) dendritic inputs, differences shown in *F*. Temporal precision is most affected at stimulus frequencies at and above CF. Yellow contours represent VS = 0.6. Presentation as in *A-C. G,H,I:* Contour plot of the mean phase φ of output AP without (*G*) and with (*H*) dendritic inputs, differences shown in *I*. Presentation as in *A-C*. Note that the phase advance mostly accompanied the changes in VS, not the changes in output AP rate.

We concluded that the interaction of the main endbulb input with phase-locked dendritic inputs shaped the output rate and precision of the model SBC in a complex, frequency-dependent manner. At CF and for frequencies below that, the number of well-timed output AP the SBC model generates was clearly increased. This could have a significant effect on temporal processing in binaural target areas of SBC in the auditory brainstem. The benefit of dendritic inputs for temporal precision was strongest for higher frequencies. In these stimulus conditions the temporal precision of the SBC response, both in the model and in real recordings (Kuenzel et al., 2011), started to deteriorate. Thus, in the model with a single endbulb and no inhibitory inputs, the dendritic inputs ameliorated the deterioration of temporal coding for higher frequencies.

### Influence of dendritic properties on the efficacy of dendritic inputs to SBC

In the following section we wanted to investigate the influence of morphological and physiological parameters of the SBC dendrite on the effect the interaction of dendritic and main endbulb inputs had. For this we again simulated responses with and without dendritic inputs similar to the experiment shown in Fig. 7. In the following however, we only show the difference (cf. Fig. 7C,F,I) in output rate, VS and preferred phase between the two conditions plotted against the parameters we varied.

We first varied morphological parameters of the SBC model dendrite (Fig. 8A1). It became evident, that best increase in output rates was achieved for dendrites above 3.3µm diameter and between 50 and 200 µm. With increasing dendrite diameters, primary dendrites had to be shorter to achieve comparable improvement in output spike rates. In fact, the highest difference in output rate between the conditions without and with dendritic inputs (52 Hz) was achieved for a dendrite of L = 81 µm and D = 6.2 µm. However, a variety of parameter combinations in the range of 3.3 µm < D < 5.5 µm and 50 µm < L < 200 µm caused robust increase in output spiking. Best improvements of temporal precision were generally found for short dendrites below L = 100 µm (Fig. 8A2). Although the best improvement of vector strength was found for a dendrite of L =33 µm and D = 9.1 µm, the improvement of temporal precision was only weakly dependent on dendrite diameter for short dendrites. In a range of diameters between 2.7 µm < D < 6 µm a robust improvement of phase-locking precision was observed for dendrites below L = 100 µm. Inputs to dendrites that were narrower or wider than this range could also improve temporal precision, however in these cases the length of the dendrite was more restrictive. Interestingly, a distinct subset of morphological parameters emerged where dendritic inputs actually caused reduction of phase-locking precision. Inputs to dendrites of L =291 µm and D = 5.3 µm reduced the vector strength by −0.12. The results for the phase advance (Fig. 8A3) largely followed those of the improvement of vector strength. Here again, inputs to a short, thick dendrite of L = 113 µm and D = 7.3 µm caused the largest phase advance of −0.11 cycles. We conclude, that morphological parameters well within the estimated physiological range for bushy cell dendrites allowed dendritic inputs to cause a robust increase in output spiking. However, the effect of improvement of phase-locking precision was limited to short dendrites below about 100µm.

**Fig. 8.**
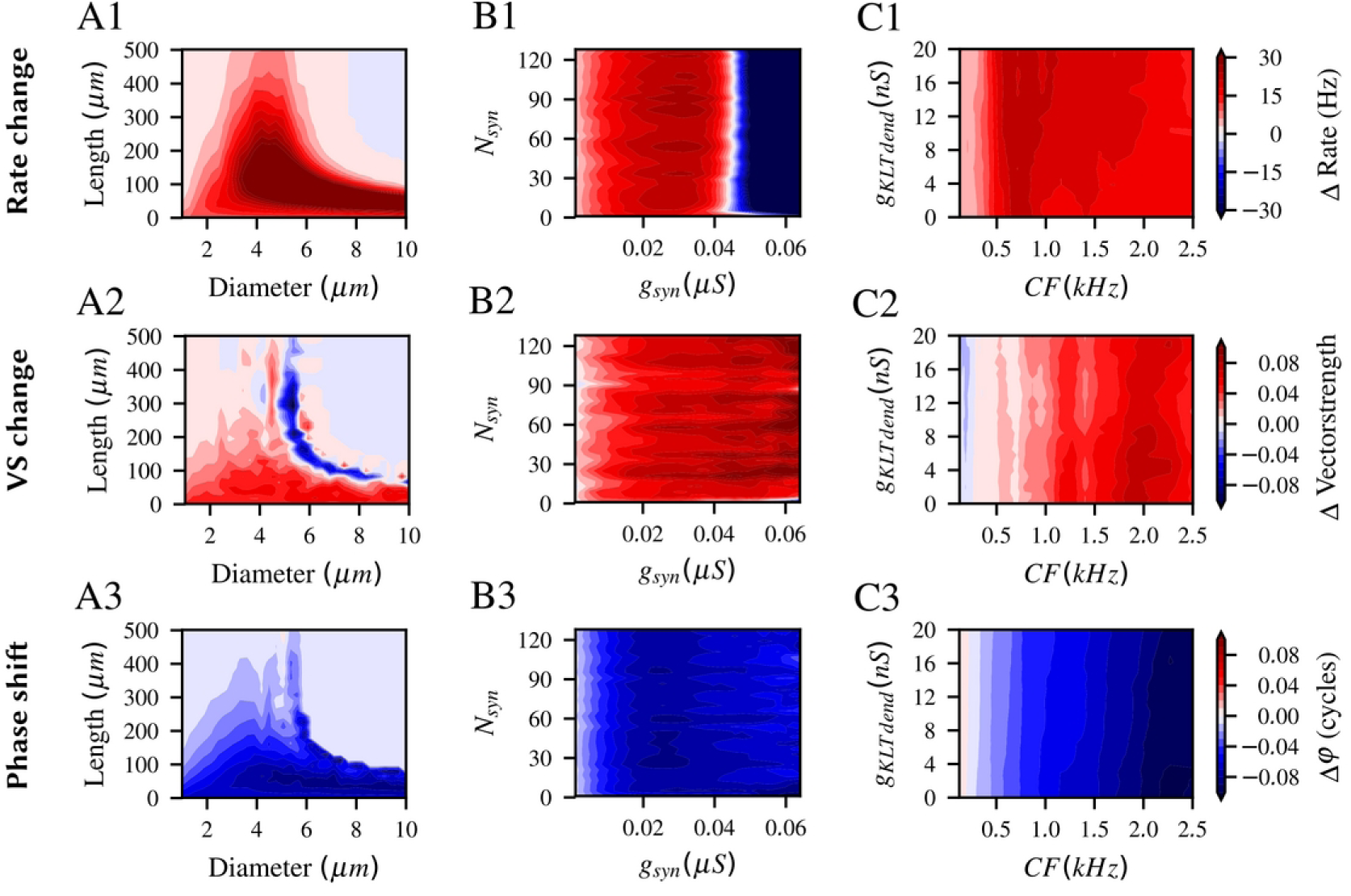
Dendritic morphology and biophysics influence the effects of dendritic inputs on output spike rate and temporal coding. *A:* Contour plots of the difference in output AP rate (*A1*), vector strength (*A2*) and preferred phase φ (*A3*) between the model without and with dendritic inputs for 1024 (32 × 32) combinations of length and diameter of the primary dendrite. Presentation of these difference plots as Fig. 7C. *B:* Contour plots of the difference in output AP rate (*B1*), vector strength (*B2*) and preferred phase φ (*B3*) between the model without and with dendritic inputs for 1024 (32 × 32) combinations of total dendritic synaptic conductance g_syn_ and the number of dendritic synaptic inputs N_syn_. Presentation of these difference plots as Fig. 7C. *C:* Contour plots of the difference in output AP rate (*C1*), vector strength (*C2*) and preferred phase φ (*C3*) between the model without and with dendritic inputs for 1024 (32 × 32) combinations of total characteristic frequency CF in kHz and the dendritic low voltage-activated potassium conductance g_KLT_. Presentation of these difference plots as Fig. 7C.

We next tested how the number and total conductance of dendritic synapses influenced the interaction of main endbulb and dendritic inputs (Fig. 8B). As expected from the subthreshold analysis (Fig. 3) the number N of dendritic synapses did not effect the outcome of the simulation for N > 10 inputs. Instead, the result predominantly depended on the total dendritic conductance. For the improvement of output rate (Fig. 8B1) an optimal range of conductance was evident: The best improvement was seen for N = 91 synapses with a total conductance of g_syn_ = 30 nS. However, above a conductance of gsyn = 44.7 ± 0.7 nS (average for all conditions with N > 3 inputs) the effect started to reverse. Here, output spike rate was actually decreased by the interaction with dendritic inputs, to a point of massive rate reduction (−155 Hz for N = 115 synapses with g_syn_ = 64nS). The temporal precision monotonically increased with increasing g_syn_ (Fig. 8B2) and was robust over a very wide range of synaptic conductances. The best (significant) vector strength improvement was found for N = 82 inputs with g_syn_ = 64 nS. Thus the few remaining output spikes in conditions of very high dendritic conductance were best locked to the stimulus phase. However, it must be noted that for these experiments at least 5 seconds of spiking activity was simulated per condition and therefore even in conditions of massive rate reduction a sufficient number of well-timed AP could easily be collected. This is most likely not a feasible mode of phase encoding in the brain. We thus concluded that we should reject the conditions that caused massive rate reductions as unrealistic, even if best improvement of temporal precision resulted from these. This notion is supported by the quantification of the phase advance (Fig. 8B3), which was maximal only for a more limited range of total dendritic conductances between roughly 20 nS and 35 nS. The conclusion from the experiments shown in Fig. 8B was, that the actual number of dendritic inputs did not matter for the beneficial effects these inputs have in the interaction with the main endbulb input, beyond a very low number at least. Furthermore, the total dendritic synaptic conductance had to be below 35 nS to generate effects that did not overtly divert from realistic behavior of SBC.

We wanted to investigate the hypothesis that dendritic g_KLT_ might tune the effect of interaction of dendritic inputs with the main endbulb to a specific range of characteristic input frequencies (cf. Fig. 5). For this we varied the dendritic low-threshold potassium conductance g_KLT_ and the CF of the model and observed effects for stimulation at CF (Fig. 8C). Improvement of output rate upon stimulation at CF (Fig. 8C1) appeared mostly to depend on CF. Highest increases were found for CFs between 500 Hz and 1000 Hz (best: +23 Hz output rate at 647 Hz, 18 nS g_KLTdend_). Nevertheless output rate increased robustly over all CFs tested, except below 250 Hz. However, contours on the flanks of the area of best improvement were skewed. Hence, output rate improvements in this area both depended on CF and, to a lesser degree, also on g_KLTdend_. Thus, in order to attain a consistent effect on output rate over a range of different CFs, g_KLTdend_ had to increase with increasing CF (on the high-frequency side). As an example we quantified here the slope for the +18 Hz output rate contour by linear regression: on the high frequency side, g_KLTdend_ needed to increase by 0.022 nS / Hz CF for a constant output rate improvement. On the low-frequency side (below 500 Hz) contours were skewed in the other direction, thus g_KLTdend_ needed to decrease with increasing CF at a steep slope of −0.19 nS / Hz CF for a constant output rate improvement. Results were comparable for temporal precision (Fig. 8C2,3). Indeed, improvement of vector strength and the accompanying phase advance mostly depended on CF. At CFs below about 750 Hz increase in VS was low or non-existant but was robustly present above this CF range. Again, an area of best improvement of VS could be identified for higher CFs between 1750 and 2250 Hz (Fig. 8C2), where phase advance was especially pronounced (Fig. 8C3). As was the case for the output rate improvement, contours of equal VS improvement and phase advance are skewed for higher CFs (Fig. 8C2,3). Thus, g_KLTdend_ in these cases needed to increase at 0.07 nS / Hz CF (quantifed for the +0.06 VS contour as an example) in order to maintain a constant improvement of VS or phase advance. Our model results therefore supported the hypothesis that dendritic g_KLT_ could indeed act as a parameter to fine-tune effects of the interaction of dendritic and main endbulb inputs in SBC to a specific range of characteristic input frequencies.

In order to quantify the frequency-dependency of the effects of the interaction of dendritic and axosomatic inputs we varied both CF and input frequencies (Fig. 9A). As was already evident from the frequency response area of a CF = 1500 Hz unit we showed in Fig. 7, best improvement of output rate was predominantly found at frequencies below CF (Fig. 9A1). Indeed, contours of equal output rate improvement are skewed diagonally. As an example we quantified the slope of the ΔAP = 18 AP/s contour by linear regression. We found a 0.58 kHz/kHz slope and an y-intercept of 0.35 kHz (y = 0.58x+0.35). This means, that for CF below about 750 Hz the frequency of best improvement was largely at CF. For higher CF the frequency of best improvement increasingly moved away from CF into the low-frequency tail. Although the highest improvement of AP output rate was found in the low frequency area (587 Hz CF, 587 Hz input frequeny, ΔAP = 25.7 AP/s) improvement of output rate was robust for a wide range of CF and input frequencies. Improvement of temporal precision (and the accompanying phase advance) followed a different pattern of frequency-dependency (Fig. 9A2,3). Regardless of CF, no increase of vector strength (and accordingly, very little phase advance) occurred for input frequencies below 500 Hz. For higher input frequencies above about 700 Hz units of all CF showed robust increase of vector strength. This was of course limited by the high-frequency flank in the tuning of the auditory nerve fibers for units of low CF. Beyond that, no further clear pattern of frequency dependency emerged. Maximal improvement of temporal precision (and phase advance) seemed to occur at CF and input frequencies of around 2000 Hz and above (max VS improvement: +0.07 for CF = 1872 Hz, 2271 Hz input frequency). Overall we concluded, that the effects on output rate and temporal precision had different but overlapping patterns of frequency-dependence.

**Fig. 9.**
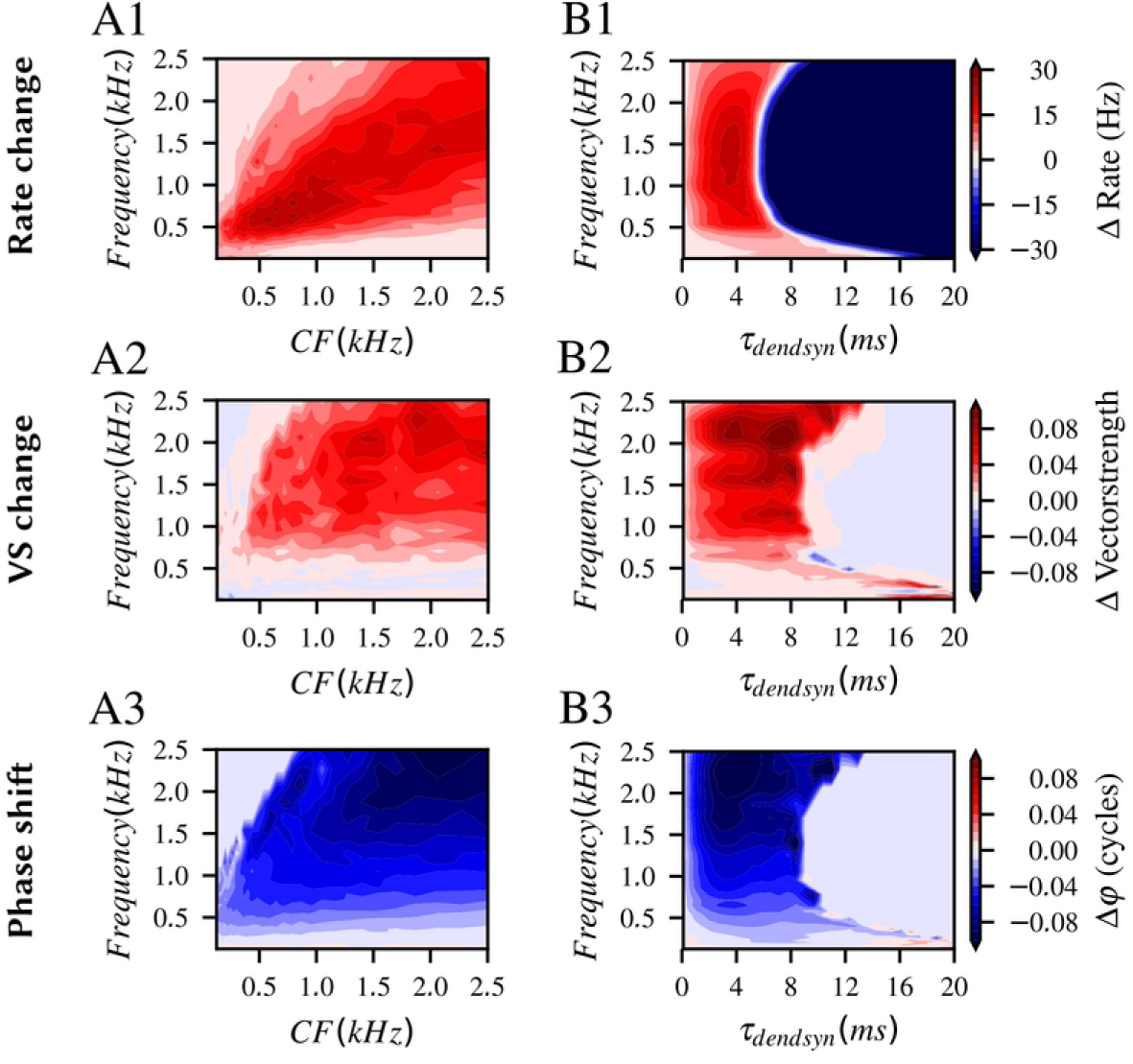
Effects of dendritic inputs on output spike rate and temporal coding depends on tonotopy and the interplay between inputs and kinetics of the dendritic synapses. *A:* Contour plots of the difference in output AP rate (*A1*), vector strength (*A2*) and preferred phase φ (*A3*) between the model without and with dendritic inputs for 1024 (32 × 32) combinations of input frequency (in kHz) and characteristic frequency of the SBC model (in kHz). Presentation of these difference plots as Fig. 7C. *B:* Contour plots of the difference in output AP rate (*B1*), vector strength (*B2*) and preferred phase φ (*B3*) between the model without and with dendritic inputs for 1024 (32 × 32) combinations of input frequency (in kHz) and the decay time-constant of dendritic synaptic inputs τ_dendsyn_. Presentation of these difference plots as Fig. 7C.

Last we hypothesized, that the frequency-dependence of the effects of the interaction of dendritic and axosomatic inputs might be governed by the kinetics of the dendritic synaptic inputs in this model. We assumed, that these synapses should have rapid kinetics, however for very low or very high input frequencies slower or even more rapid kinetics might be favorable. We therefore varied the decay time-constant of the dendritic synapse model (τ_dendsyn_) and the input frequency. CF remained at 1500 Hz in this experiment (Fig. 9B). Output rate improvement (Fig. 9B1) mostly depended on τ_dendsyn_ in this experiment. It was robust for dendritic synaptic time-constants between 0.8 ms and 6 ms. Best increase of output rate occurred at τ_dendsyn_ = 3.3 ms and 1272 Hz input frequency. A very sharp cutoff of efficacy in increasing the output rate occured above τ_dendsyn_ = 6 ms for input frequencies around CF. Higher dendritic time-constants were very detrimental for output spiking: Summation of dendritic EPSP here caused a tonic depolarization sufficient to strongly reduce the excitability of the SBC model. Interestingly, this cutoff value shows some frequency-dependency insofar as it extents towards much longer τ_dendsyn_ for frequencies below about 500 Hz, reaching 14 ms for the lowest frequencies tested (125 Hz). Surprisingly, the cutoff value also extents towards longer time-constants for frequencies above 2000 Hz. A very similar pattern was evident for the quantification of the increase of temporal precision (Fig. 9B2) and advance of the preferred phase (Fig. 9B3). However, robust improvement of vector strength and advance of preferred phase occurred over a somewhat wider range of τ_dendsyn_ values. Accordingly, the cutoff of the effect for input frequencies around CF occurred at higher τ_dendsyn_ of 9 ms. This can be understood, since the effect on temporal precision and preferred phase mostly relied on the rising slopes of the V_m_ modulation (Fig. 6F&G) and even for very long dendritic decay-time constants detectable V_m_ modulation did occur (Fig. 4A2,B2). In contrast to this, the bulk of the effect of the increase in AP output rate seemed to be carried by the V_m_ depolarization, as it tonically moved the V_m_ closer to AP threshold. The tonic depolarization however monotonically rose with τ_dendsyn_ (Fig. 4A1) and at some point, as we showed here, reached values at which the positive effects were countered by the reduction of excitability (increased Na_v_ inactivation and KLT activation). Taken together, we concluded that phase-locked synaptic inputs with rather rapid synaptic time-constants of decay are a necessity to robustly exploit the effects of the dendritic synaptic inputs for SBC coding.

Overall our experiments in this section showed that the morphological and physiological parameters of the SBC dendrite and the synaptic inputs that connect there had considerable influence on the efficacy of the dendritic inputs in supporting the main endbulb input. Some dendritic features (length, diameter, expression of ionic conductances) might indeed serve as plausible biological tuning parameters to finely adjust the effect of dendritic inputs for specific input frequencies along the tonotopic axis or divergent coding demands of subpopulations of SBC.

### Simulated dendritic inputs to realistic bushy dendrites

Up to this point all conclusions were drawn from simulations that only included a stylized dendritic model with a linear arrangement of synaptic inputs. In the next step we wanted to test whether these conclusion also hold true in a complex, 3-dimensional branched dendrite model with randomly placed synapses that is closer to the real shape of SBC in the brain. We thus reconstructed dendrites from confocal stacks of gerbil SBC, which were filled with biocytin during whole-cell patch recordings (Fig. 10). We analyzed a total of N = 18 cells which had short dendrites that reached on average only up to 83 ± 19 µm from the soma. However, the dendrites were dense and quite complex with a mean number of 36 ± 16 branchpoints and a total length of dendritic segments of 1463 ± 667 µm. A Sholl-analysis (Sholl, 1953) in 3D-space revealed a critical value of 18 ± 3 intersects at a critical range of 38 ± 12 µm. Mean value of intersects over all radii was 8± 2 intersects. These morphological data emphasized, that SBC dendrites did not reach far away from the location of the soma but showed surprisingly high complexity. The critical range could be interpreted as the distance from the soma, at which the cell receives most inputs with its dendritic tree. This was in our morphological reconstructions roughly 2 soma-diameters away from the cell under consideration and was thus most probably an area of the same or very similar characteristic frequency range of AN inputs. This illustrated well how locally dense the network of SBC dendrites in the gerbil AVCN must be (Gómez-Nieto and Rubio, 2009). The 3D-reconstructed (and volume fitted – see method section) data for initially three SBC was carefully checked for consistency and imported as segments in NEURON. We attached the dendritic part to the stylized soma and axon segments of our SBC model. This was done to compare only the properties of the realistic dendritic tree, i.e. soma and axon compartments (white and red segments in Fig. 10B&C) were discarded for the following initial analysis. When we simulated phase-locked dendritic inputs and the interaction of their effects with the main axosomatic endbulb inputs we found qualitatively similar results as with the stylized dendrite (Fig. 11). However, individual dendritic synapses were electrotonically much further from the soma and thus much less effective. Thus, higher total dendritic conductances were needed to also achieve quantitatively comparable effects. This is illustrated in Fig. 11, were we simulated frequency response areas of a SBC with realistic dendritic tree without (g_dendsyn_ = 0 nS) and with dendritic inputs, identical to Fig. 7. The overall features of the interaction of dendritic and endbulb inputs were identical: best increase of output rate for frequencies below CF (Fig. 11A-C), best increase of temporal precision (Fig. 11D-F) and strongest phase advance (Fig. 11G-I) for frequencies >1500 Hz. In order to achieve these effects a higher total dendritic synaptic conductance (g_dendsyn_ = 64 nS) was used. This can be understood as a greater electrotonic distance to the soma of a given dendritic synapse in the 3D-dendrite compared to the stylized dendrites. Of course the total membrane area, and thus the leak conductance, of the reconstructed dendrite is higher than the stylized dendrite. The efficacy of the dendritic synapses was also strongly influenced by narrow segments in the reconstructed structure, even when we enforced a minimal diameter of 1µm. No qualitative differences between the different dendritic models tested were seen.

**Fig. 10.**
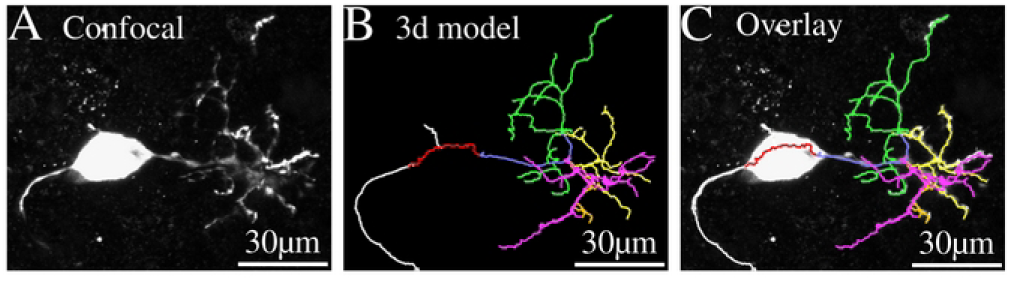
3D representation of the bushy dendrite for the SBC compartment model. *A:* Confocal image (maximal projection) of a gerbil SBC filled with biocytin during whole-cell path recording in acute brain slices in vitro. *B:* 3D model of the dendritic segments reconstructed from the confocal images in *A*. Soma (red) and axon sections (white) visible in the left half of the image, were not used for modeling, only the proximal dendritic section (blue) and distal dendritic arbors attached to this (yellow, green, magenta) were used for the compartment model.

**Fig. 11.**
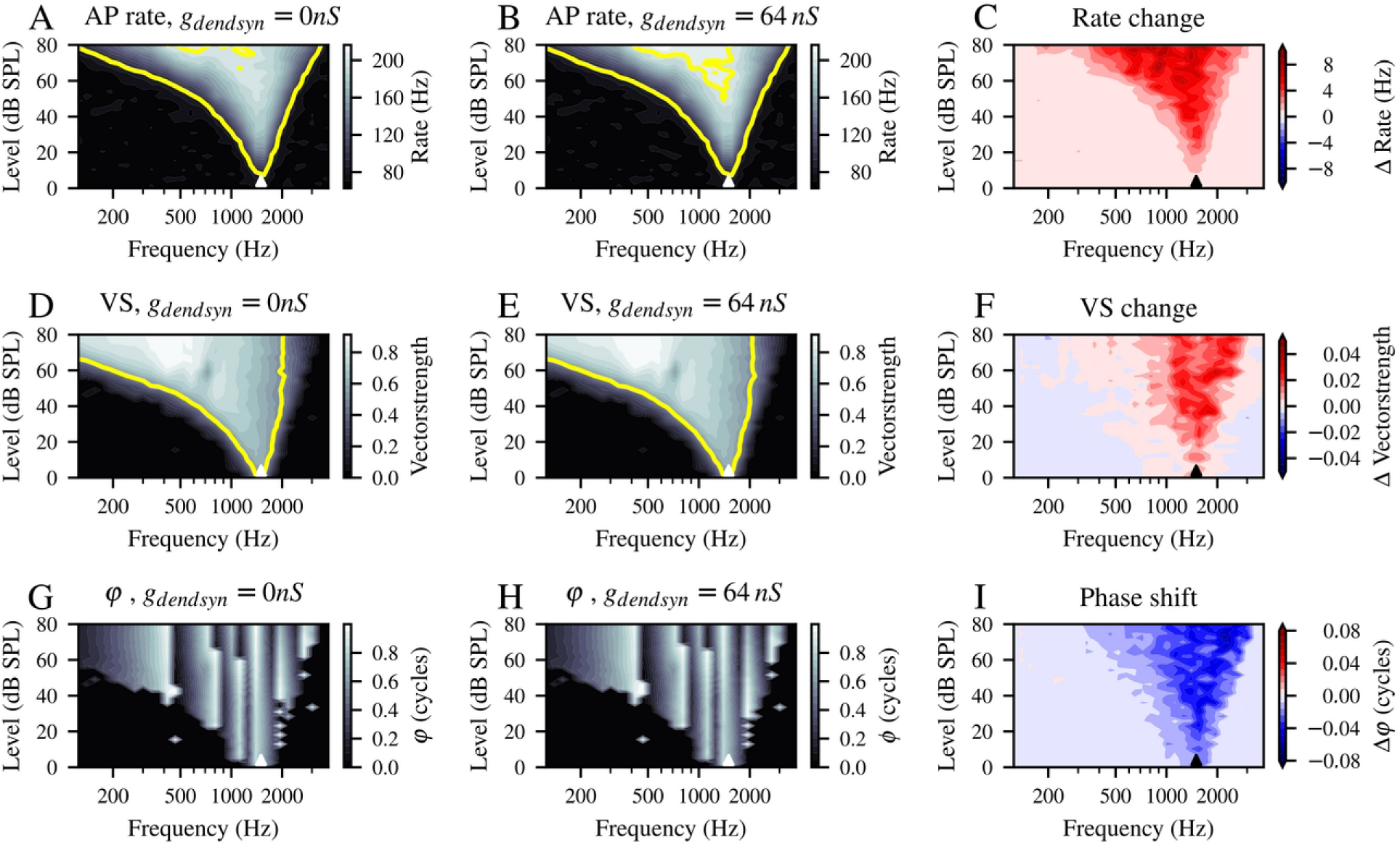
Effect of dendritic inputs connected to realistic 3D dendritic structures are qualitatively identical to effects seen with the stylized dendrite model. *A-I:* Contour plots of the output AP rate (*A-C*), temporal precision (*D-F*) and preferred phase (*G-I*) of the SBC model with a 3D dendrite model. Stimulus conditions and data presentation identical to Fig. 7.

In summary we take these results obtained with realistic 3D dendrites as first evidence that our conclusions drawn from the stylized dendritic model are also valid for complex dendritic structures as described for SBC in the brain.

## Discussion

In this study we demonstrated with a biologically plausible compartment model of spherical bushy cells that small, phase-locked dendritic excitatory inputs can augment temporal coding. The phase-locked activity of the dendritic inputs caused both tonic excitation and a modulation of the RMP at the frequency of the input. Interaction of a single axosomatic endbulb of Held input with the rising phase of the RMP modulation enhanced the efficacy of the axosomatic input in a phase-dependent manner. Together with the tonic excitation this effect caused an increase in the number and temporal precision of output spikes of the model SBC.

For SBC that receive only a single axosomatic input we concluded, that phase-locked dendritic inputs are a plausible mechanism to improve temporal precision. The postsynaptic acitivity of dendritic inputs almost certainly would be inconspicuous in single-unit recordings in vivo or may be misclassified as microphonic responses of AN fibers. We hesitate to call this a novel suggestion, as this was one of the ideas brought up before in the context of the spherical cell puzzle (Joris and Smith, 2008). However, unlike the case of many interacting subthreshold inputs (Rothman et al., 1993; Xu-Friedman and Regehr, 2005a) and few suprathreshold inputs (Xu-Friedman and Regehr, 2005b) this arrangement of input strengths had not yet been theoretically explored before in the context of temporal coding. Our study thus represents an additional step towards understanding the remarkable complexity (Kuenzel, 2019) of this station of the auditory pathway.

The improvement in temporal precision, quantified as the difference in output vector strength without and with dendritic inputs (Fig. 7) was robust but quantitatively moderate. One could thus argue, that dendritic inputs only really play a minor role in temporal coding. In the same type of SBC model quantitatively similar findings were made for the effect of inhibitory inputs (Nerlich et al., 2014; Kuenzel et al., 2015) and cholinergic excitation (Goyer et al., 2016) on temporal precision. It is very hard to judge however, how these parallel mechanisms would interact in the intact brain during natural listening situations. We hypothesize that the combined contribution of individually smaller effects could summate substantially as well as provide dynamic flexibility for different coding demands. Furthermore, given the strong focus on temporal precision in sound localization circuitry of the medial superior olive (van der Heijden et al., 2013; Franken et al., 2014; Plauška et al., 2016) and the relatively low amount of convergence present at this station (Couchman et al., 2010), small increases in rate and precision of bushy cell outputs could have considerable functional impacts for sound localization using interaural time differences.

When we explored the influence of biophysical and morphological parameters on the effect of dendritic inputs (Figs. 1,5,8 & 9) our results suggested that dendritic parameters could be tuned to emphasize a specific effect (tonic vs. modulated excitation) and or matched to a specific range of input frequencies. The latter would argue for gradients of dendritic properties along the tonotopic axis of the ventral cochlear nucleus. For the avian homologue of the VCN, the nucleus magnocellularis, it was established that gradients of biophysical parameters (Fukui and Ohmori, 2004; Oline et al., 2016; Hong et al., 2018), especially voltage activated potassium conductances, optimize temporal coding for a given range of input frequencies. Furthermore, dendritic morphology of neurons in the chicken NM changes drastically with tonotopic position: low frequency neurons (Wang et al., 2017) have long dendrites while high frequency neurons are essentially adendritic. Tonotopic gradients are even more pronounced in nucleus laminaris (the analogue of the medial superior olive). Here both biophysical parameters (Kuba et al., 2005) and dendritic length (Korn et al., 2011) smoothly varied with tonotopic position and it was convincingly shown that dendritic filters are tuned to match the tonotopic position (Slee et al., 2010). We thus think that it is not unreasonable to assume that similar tonotopic gradients could be present in the mammalian VCN. Unfortunately, to our knowledge, no study systematically analyzed biophysical nor morphological parameters along the tonotopic axis in the mammalian VCN. The oblique orientation of the tonotopic axis in the mammalian VCN (Ryugo and Parks, 2003; Muniak et al., 2013) is the most likely reason for that. While Lauer et al. (2013) indeed show compelling ultrastructural and morphological differences between anterior and posterior positions they do not reconstruct or quantify dendritic structures. In the VCN, specifically for bushy cells, cell size does roughly correlate with tonotopic position as low frequency BC, if present, are larger than high frequency BC (Cant and Casseday, 1986; Bazwinsky et al., 2008) (cat; gerbil). However, no systematic differences of dendritic shapes have been reported or analyzed yet. Given the complex and dense nature of the bushy dendrites it might well be that simple morphological metrics (long vs. short path, narrow vs. wide field) do not catch the essential tuning parameters. A careful 3D-reconstruction of numerous bushy dendrites, as we (in this study) and others have begun, will in our opinion be necessary to resolve this question. Unfortunately at this moment our set of dendritic reconstruction derived from in vitro patch recordings in oblique brain slices, only offers rough positional estimates on the tonotopic identity of the cells. A systematic study of spherical bushy cell dendritic morphology filled in-vivo after single-unit recordings (Pinault, 1996; Kuenzel et al., 2011) would be most useful for this question. Nevertheless, from binaural nuclei of the mammalian auditory brainstem it is well known, that cell size (Weatherstone et al., 2017) (MNTB) and biophysical properties (Barnes-Davies et al., 2004; von Hehn et al., 2004; Leao et al., 2006) (LSO; both MNTB) can vary with tonotopic position. We overall conclude that tuning of postsynaptic and dendritic parameters along the tonotopic axis also in the mammalian VCN is a plausible proposal.

Since very little is known about the properties of bushy dendrites and dendritic synaptic inputs, one wonders how biologically plausible our parameters were. First, the fact that at least some of these inputs are axodendritic contacts of endbulb structures contacting another SBC soma (Ostapoff and Morest, 1991; Ryugo and Sento, 1991; Gómez-Nieto and Rubio, 2009) lead us to model these dendritic inputs with rapid kinetics and robust temporal precision. The actual number of inputs does seem much less critical than the total synaptic conductance (Fig. 8B1), as higher total conductances could cause erroneous spiking (Fig. 3E3) or strong reduction in output rates (Fig. 8B1). To us this demonstrates the use of our in-silico approach, as unknown parameters can be at least narrowed to a more biologically plausible range. Other dendritic excitatory inputs (Kuenzel, 2019) to bushy cells do not necessarily have robust phase-locking or rapid kinetics. Our results show that these inputs, given the total active dendritic conductance remains in a reasonable range, would still contribute to temporal processing by increasing the amount of well-timed spikes the cell generates upon endbulb of Held input activity. It was recently shown that even non-auditory inputs to the VCN can influence temporal coding (Heeringa et al., 2018). Our model also provides mechanistic explanations for these findings and suggests a common role of excitatory inputs to spherical bushy cell dendrites: dendritic inputs gate and modulate the processing of auditory temporal information along the main endbulb-soma-axon path.

Finally we want to comment on the realistic 3D-dendritic structures we used in a first attempt to understand the role of the bushy dendritic structure. While we have confindence in the reconstruction of the overall shape and connectivity of dendrites, the faithful estimation of the diameter of small dendritic segments based on confocal laser-scanning microscopy was difficult due to technical limitations. Since the algorithm used produced very narrow segments (< 0.3 µm) whenever the fluorescent signal was locally weak, we enforced a minimal diameter of 1 µm. Still, rapid changes of narrow dendritic segments and local swellings of larger diameter were commonly seen. Whether this is an artefact of the fixation and staining or biologically relevant we can not state at this moment. We thus refrained from starting a more in depth analysis of the properties of the 3D-dendrites (i.e. mapping electrotonic distance) at this point. The technical difficulties with volume estimation are a known limitation in reconstructing 3D dendritic structures (Ascoli et al., 2001) and an area of active research (Ming et al., 2013; Luo et al., 2015). In the future more sophisticated methods should be employed to derive segment diameters from 3D microscopy data. Indeed, a morphological dataset of tonotopically identified spherical bushy cells obtained with super- or ultra-resolution (Holcomb et al., 2013) methods would greatly help modeling efforts to better understand the peculiar nature of the bushy dendritic structure of this cell type. Nevertheless we are convinced that our modeling effort provides interesting insights into the dendritic function at this level of the auditory pathway and opens promising avenues for further experimental work in this system.

## Conflict of Interest Statement

The authors declare that the research was conducted in the absence of any commercial or financial relationships that could be construed as a potential conflict of interest.

## Acknowledgments

We thank Stefanie Kurth and Charlène Gillet (Department of Chemosensation, Institute of Biology II, RWTH Aachen University) for providing biocytin-labeled bushy cells and excellent technical assistance with the 3D-reconstruction of neurons.

## Notes

Funding: This work was supported by the DFG Priority Program 1608 “Ultrafast and temporally precise information processing: Normal and dysfunctional hearing” [KU2529/2-2]

### Competing Interest Statement

The authors have declared no competing interest.

